# A collection of programs for one-dimensional Ising analysis of linear repeat proteins with point substitutions

**DOI:** 10.1101/2020.06.27.175224

**Authors:** Jacob D. Marold, Kevin Sforza, Kathryn Geiger-Schuller, Tural Aksel, Sean Klein, Mark Petersen, Ekaterina Poliakova-Georgantas, Doug Barrick

## Abstract

A collection of programs is presented to analyze the thermodynamics of folding of linear repeat proteins using a 1D Ising model to determine intrinsic folding and interfacial coupling free energies. Expressions for folding transitions are generated for a series of constructs with different repeat numbers and are globally fitted to transitions for these constructs. These programs are designed to analyze Ising parameters for capped homopolymeric consensus repeat constructs as well as heteropolymeric constructs that contain point substitutions, providing a rigorous framework for analysis of the effects of mutation on intrinsic and directional (i.e., N- versus C-terminal) interfacial coupling free-energies. A bootstrap analysis is provided to estimate parameter uncertainty as well as correlations among fitted parameters. Rigorous statistical analysis is essential for interpreting fits using the complex models required for Ising analysis of repeat proteins, especially heteropolymeric repeat proteins. Programs described here are available at https://github.com/barricklab-at-jhu/Ising_programs.

## 1. INTRODUCTION

One of the goals of protein folding studies is to quantify the contributions of specific structural features within the native state to the overall free energy of folding. As a result of the high level of cooperativity in protein folding, the relative contributions of such structural features cannot easily be determined because they are hidden in an all-or-none, or “two-state” transition. Though two-state behavior makes overall protein folding energetics easy to quantify, it prevents an energetic dissection of the whole into its parts.

Over the last 15 years, a class of proteins with rough translational symmetry, termed “linear repeat proteins”, has been recognized as having an architecture that permits quantification of local stabilities, long-range coupling energies, and cooperativity (Mello and Barrick, 2004; Kajander et al., 2005; Wetzel et al., 2008; Aksel et al., 2011; Marold et al., 2015; Geiger-Schuller and Barrick, 2016). Much like the helix-coil transitions of simple polypeptides, the unfolding of linear repeat proteins can be analyzed using a one-dimensional Ising model, where the overall folding free energy can be broken down into intrinsic folding energies of individual repeats (*ΔG*_*i*_) and coupling energies between folded nearest-neighbor repeats (*ΔG*_*i−1,i*_).

To quantify cooperativity in linear repeat proteins, a series of equilibrium unfolding transitions (often from chemical denaturation) are obtained for proteins containing different numbers of repeats, and the transitions are globally fitted using a model that relates the extent of folding to *ΔG*_*i*_ and *ΔG*_*i−1,i*_. Varying the number of repeats is essential to resolve the values of *ΔG*_*i*_ and *ΔG*_*i−1,i*_ (Aksel and Barrick, 2009). Although this analysis is simplest when applied to arrays with identical repeat sequence, solubility considerations usually require sequence modification to one or both terminal repeats (Figure 1A). The effects of these terminal modifications on stability need to be accounted for by including intrinsic folding energy parameters for the substituted terminal repeat (*ΔG*_*N*_ and *ΔG*_*C*_).

**Figure 1.**
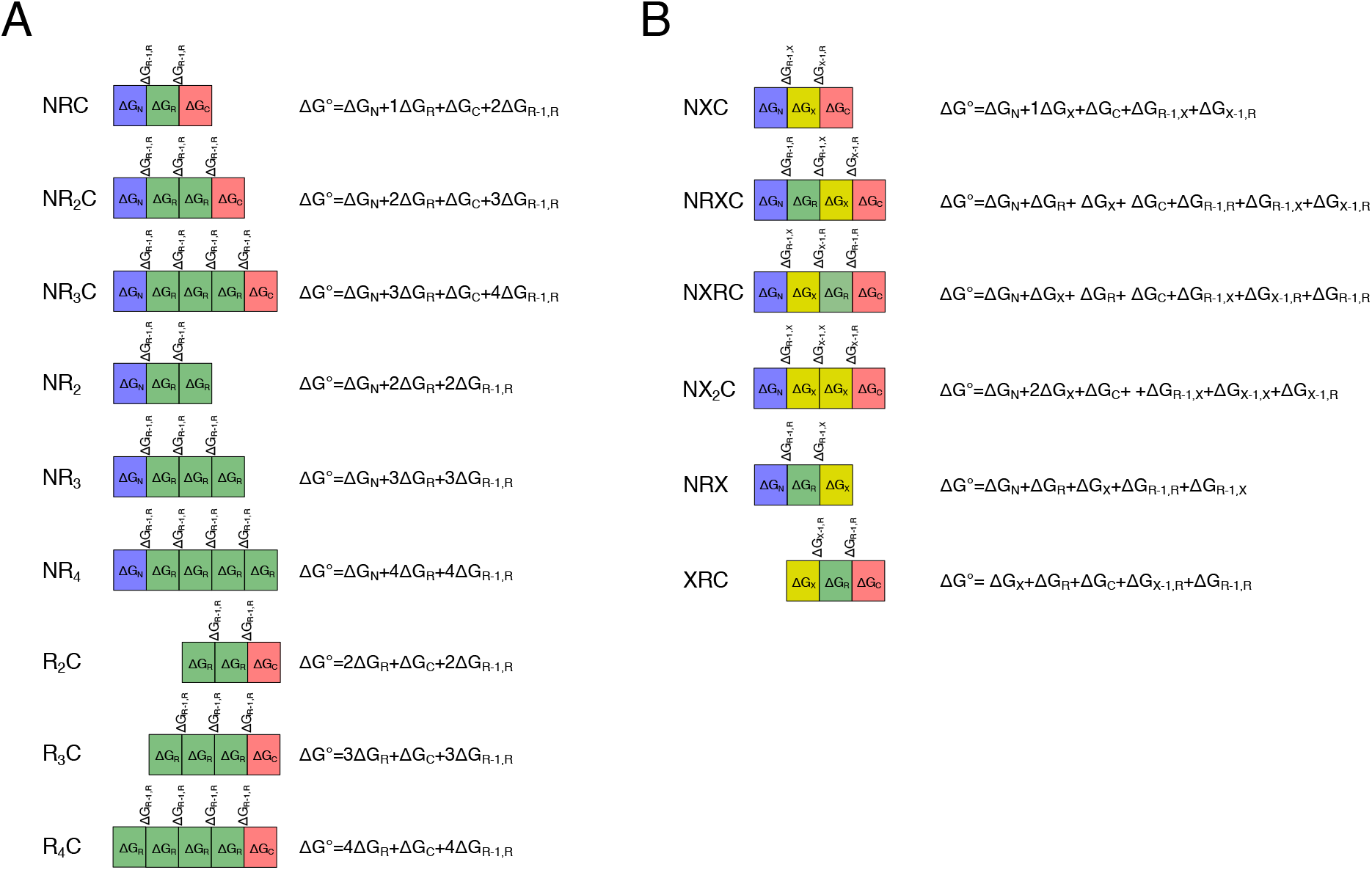
Schematic of a series of tandem repeat constructs for Ising analysis. Each repeat is indicated by a rectangle. (A) A capped homopolymeric series. Identical repeats (R) are colored green; to maintain solubility, these repeats are flanked on the N- or C-terminus (or both) by polar capping repeats (N, blue and C, red, respectively). (B) A capped heteropolymeric series, in which repeats harboring a sequence substitution (X, yellow) are combined with R, N, and C repeats. Each repeat has an associated intrinsic folding energy (ΔG_i_), depending on its type (N, R, X, or C). Folded repeats couple with their neighbors through an interaction energy (ΔG_i−1,i_) that is assumed to the same for NR, RR, and RC interfaces. The equations on the right give the free energy difference between the fully folded and fully unfolded states as a sum of these intrinsic and interfacial terms. Global Ising analysis of the series in (A) allows the N, R, and C intrinsic and interfacial parameters to be determined. By combining the series in (A) and (B), the effects of mutation (X) on Ising parameters can be determined (see Table 1).

In addition, by combining point mutation and length variation, Ising analysis of linear repeat proteins can be extended to analyze the effects of sequence substitutions on the underlying parameters (Figure 1B). Although these sequence perturbations increase the complexity of the fitting model, Ising analysis of constructs containing sequence substitutions along with the capped homopolymer constructs in Figure 1A allows the energetics of specific structural features to be resolved into intrinsic and interfacial stabilities (Aksel et al., 2011; Cortajarena et al., 2011).

**Table 1.**
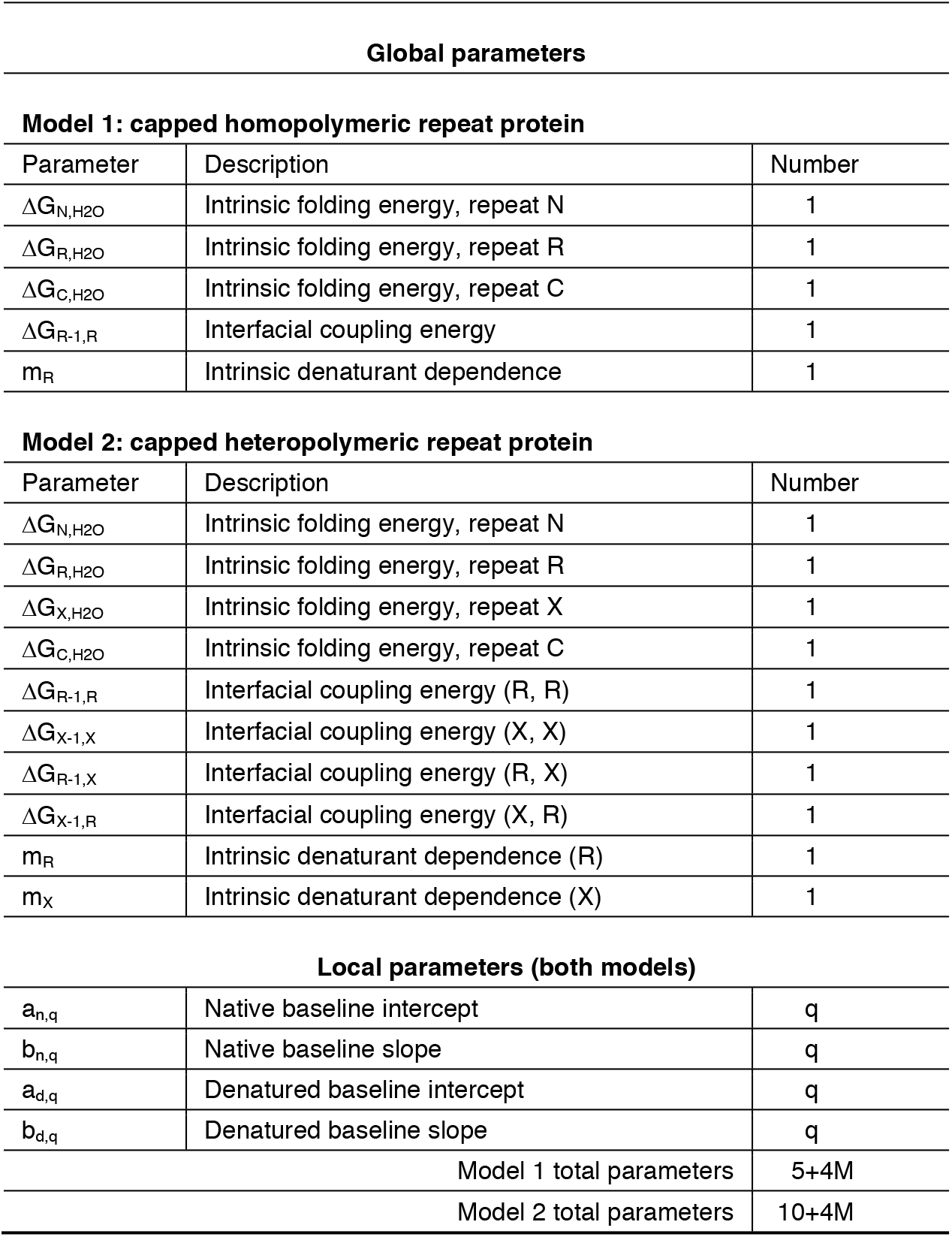
Adjustable parameters for capped homopolymer and heteropolymer models.

| Global parameters |  |  |
| --- | --- | --- |
| **Model 1: capped homopolymeric repeat protein** |  |  |
| Parameter | Description | Number |
| $\Delta G_{N,H_2O}$ | Intrinsic folding energy, repeat N | 1 |
| $\Delta G_{R,H_2O}$ | Intrinsic folding energy, repeat R | 1 |
| $\Delta G_{C,H_2O}$ | Intrinsic folding energy, repeat C | 1 |
| $\Delta G_{R-1,R}$ | Interfacial coupling energy | 1 |
| $m_R$ | Intrinsic denaturant dependence | 1 |
| **Model 2: capped heteropolymeric repeat protein** |  |  |
| Parameter | Description | Number |
| $\Delta G_{N,H_2O}$ | Intrinsic folding energy, repeat N | 1 |
| $\Delta G_{R,H_2O}$ | Intrinsic folding energy, repeat R | 1 |
| $\Delta G_{X,H_2O}$ | Intrinsic folding energy, repeat X | 1 |
| $\Delta G_{C,H_2O}$ | Intrinsic folding energy, repeat C | 1 |
| $\Delta G_{R-1,R}$ | Interfacial coupling energy (R, R) | 1 |
| $\Delta G_{X-1,X}$ | Interfacial coupling energy (X, X) | 1 |
| $\Delta G_{R-1,X}$ | Interfacial coupling energy (R, X) | 1 |
| $\Delta G_{X-1,R}$ | Interfacial coupling energy (X, R) | 1 |
| $m_R$ | Intrinsic denaturant dependence (R) | 1 |
| $m_X$ | Intrinsic denaturant dependence (X) | 1 |
| Local parameters (both models) |  |  |
| $a_{n,q}$ | Native baseline intercept | q |
| $b_{n,q}$ | Native baseline slope | q |
| $a_{d,q}$ | Denatured baseline intercept | q |
| $b_{d,q}$ | Denatured baseline slope | q |
| Model 1 total parameters |  | 5+4M |
| Model 2 total parameters |  | 10+4M |

Here we present a collection of programs that perform 1D-Ising analysis on repeat protein arrays. The programs return fitted parameter values for *ΔG*_*i*_ and *ΔG*_*i−1,i*_, and their denaturant dependences. In addition, bootstrap analysis (Efron, 1979; Johnson, 2008) is used to determine confidence intervals for each parameter and their correlations with each other. We use these programs to analyze a series of unfolding transitions of homopolymeric repeat arrays with modified caps, and to analyze a larger data set that includes point substitutions in internal repeats. Analysis of point substitutions allows the energetics associated with atomic-level structural features to be resolved into intrinsic and coupling energies. Moreover, by creating asymmetric interfaces, the coupling contributions of these structural features can be apportioned into the N- (*i−1, i*) and C-terminal (*i, i+1*) interfaces. This suite of programs, which is available on github (https://github.com/barricklab-at-jhu/Ising_programs) as a set of python programs and Jupyter notebooks, significantly extends an earlier analysis suite of programs (Lowe et al., 2018) by providing analysis of mutational data, bootstrapped confidence intervals, and full and partial parameter correlation analysis.

## 2. REPRESENTATION OF FOLDING TRANSITIONS OF REPEAT PROTEINS USING AN ISING MODEL

To determine *ΔG*_*i*_ and *ΔG*_*i, i−1*_ from Ising analysis, equilibrium folding transitions (colloquially, “melts”) need to be acquired for a series of constructs of varying repeat number (e.g., Figure 1). These melts are most easily obtained using chemical denaturation with urea or guanidine hydrochloride. Although thermal unfolding provides access to important quantities such as enthalpy and entropy, thermal denaturation is much less likely to be reversible than chemical denaturation, preventing the determination of true equilibrium thermodynamic parameters. Folding is often monitored by circular dichroism (CD) spectroscopy, which is well suited for α-helical repeats. Other spectroscopic signals, such as intrinsic tryptophan fluorescence, may also be suitable for monitoring unfolding. Typically, multiple melts are measured for each type of construct. The resulting data set, which contains M melts for all constructs, is globally fitted to estimate Ising parameters using the equations below.

### 2.1 A matrix approach to computing partition functions for linear repeat proteins

Unlike equilibrium two-state models for protein folding, the Ising model includes intermediate states where some repeats are folded and others are unfolded. Analysis of unfolding transitions using this model requires an expression for folding that includes all 2^*ℓ*^ conformations (where *ℓ* is the number of repeats in a particular construct). Such an expression can be obtained from a partition function in which folding is represented at the level of individual repeats using two-by-two correlation matrices (*W*). For a general heteropolymer with *ℓ* repeats,

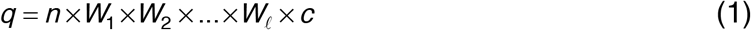

where

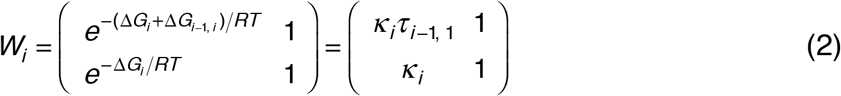

The top and bottom rows correspond to repeat *i−1* being folded and unfolded. The left and right columns correspond to repeat *i* being folded and unfolded. The quantities *κ*_*i*_ and *τ*_*i−1, i*_ are equilibrium constants for intrinsic folding of repeat *i*, and for interface formation between repeats *i−1* and *i*. The *n* and *c* terms in equation 1 are vectors that convert the *W* matrix product (itself a two-by-two matrix) into a scalar:

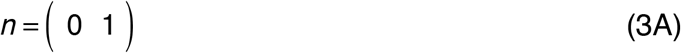

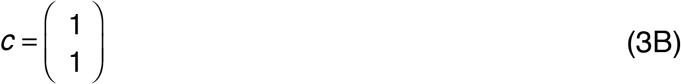

The zero in equation 3A eliminates terms that include a folded ghost repeat^1^ at position 0 (corresponding to the top row of the matrix product in equation 1).

### 2.2 Simplification of the partition function for capped homopolymeric repeat proteins

The product of the *W*_*i*_ matrices can be simplified considerably if repeats are identical. For constructs with *ℓ* − 2 identical internal R repeats and terminal N- and C-caps (see the capped homopolymeric series, Figure 1A), the partition function can be written

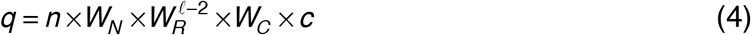

Although the same number of matrix multiplications are needed to compute the homopolymeric and the general partition function (equations 4 and 1, respectively), the homopolymeric version involves many fewer free energy terms. Here we refer to a model used to analyze capped homopolymeric repeat proteins as “model 1” (Table 1). Since the solubilizing substitutions on the capping repeats are likely to affect the intrinsic free energy, separate free energy terms (ΔG_N_, ΔG_R_, and ΔG_C_) are included in the model. However, since these solubilizing substitutions are distant from the interfaces with the internal R repeats, a single interfacial free energy is used to model all interfaces (NR, RR, and RC interfaces), which we will represent as ΔG_R−1,R_.

### 2.3 Modification of the partition function for capped heteropolymeric repeat proteins

For several decades, point substitutions have been used in protein folding studies to determine how specific interactions contribute to protein stability (Alber and Matthews, 1987) and to the kinetics of folding (Matthews and Hurle, 1987; Fersht et al., 1992).

Combining targeted sequence substitutions with Ising analysis of repeat proteins permits the effects of structural perturbation on stability to be resolved into intrinsic and interfacial free energies. One way to quantify the effects of point substitution on intrinsic and interfacial stability is to prepare and analyze capped homopolymeric sequences with the same substitution in each repeat, and compare fitted Ising parameters to those determined from unsubstituted repeats. In this approach, ΔG_R_ and ΔG_R−1,R_ values are fitted separately for each series using the capped homopolymer partition function (equation 4). Comparing these values reveals the effects of substitution on intrinsic folding and interfacial coupling. This approach was used to determine the effects of surface substitution within the helices of consensus TPR arrays (Cortajarena et al., 2011); substitutions were found to perturb ΔG_R_ (although ΔG_R−1,1_ was not varied in fits of the substituted arrays).

A more informative approach to quantify the effects of point substitution on Ising parameters involves constructing arrays in which unsubstituted and substituted repeats are combined in the same constructs (Figure 1B). In this heteropolymeric variation, the partition function contains different *W* matrices for unsubstituted versus unsubstituted repeats (which we will refer to as X and R repeats), which differ in their intrinsic folding free energy parameters (ΔG_X_ and ΔG_R_). Because the heteropolymer approach creates asymmetric interfaces (RX, XR, and XX, in addition to RR; see Figure 1B), these *W* matrices are further subdivided depending on the identity (X versus R) of their *i−1* repeat, which differ in their interfacial coupling energies (ΔG_R−1,X_, ΔG_X−1,R_, ΔG_X−1,X_, and ΔG_R−1,R_). Although this approach increases the complexity of the model (compare model 2 with model 1 in Table 1), additional structural information is accessible using this approach. Specifically, comparing ΔG_R−1,X_ and ΔG_X−1,R_ values to ΔG_R−1,R_ resolves whether the substitution in the X repeat at position *i* affects the stability of the *i−1* interface or the *i+1* interface. Moreover, comparing ΔG_X−1,X_ with the sum of ΔG_R−1,X_ and ΔG_X−1,R_ provides a measure of thermodynamic coupling between adjacent interfaces.

### 2.4. Using the partition function to model denaturant-induced unfolding transitions

Unfolding transitions can be modeled using the fraction of repeats that are folded as a function of denaturant concentration. The fraction of repeats that are folded (*f*_*f*_ = *n*_*f*_/(*n*_*f*_ + *n*_*u*_)) can be calculated by partial differentiation of the partition function with respect to intrinsic equilibrium constants of each type. For model 1 (Table 1),

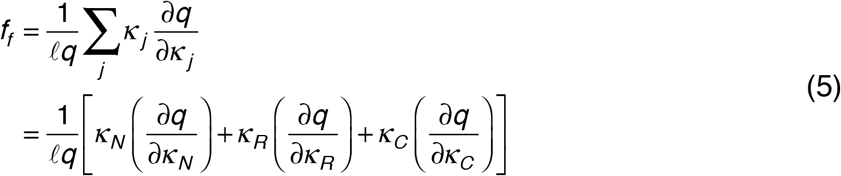

For model 2, equation 5 contains an additional partial derivative (with respect to *κ*_*x*_). Denaturant dependencies are typically built into the intrinsic (*κ*_*i*_) but not the interfacial (*τ*_*i, i−1*_) terms,^2^ assuming a linear dependence of free energy on denaturant concentration,

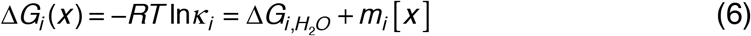

where [*x*] is the molar denaturant concentration, and *ΔG*_*i,H2O*_ and *m*_*i*_ are constants determined from the fit. The *H_2_O* subscript associated with the free energy on the right-hand side of equation 6, which indicates an extrapolated free energy in the absence of denaturant (Greene and Pace, 1974), will be omitted below for brevity. For analysis of heteropolymeric repeat arrays, a denaturant dependence for both types of repeats can be treated separately (*m*_*R*_ and *m*_*X*_; model 2).

Using *f*_*f*_, the observed signal for each of the M melts can be calculated as a population-weighted average:

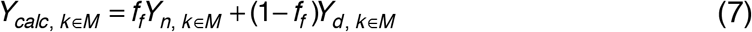

where *Y*_*n, k∈M*_ and *Y*_*d, k∈M*_ are spectroscopic signals of the fully folded and fully unfolded states (see equation 9 below) for the *k*^th^ of M melts^3^. Note that equation 7 assumes equal spectroscopic contributions of all repeats, which is likely to be the case for a signal such as CD, which measures secondary structure in each repeat. Equation 7 can be modified to account for site-specific signals such as tryptophan fluorescence (see Aksel and Barrick, 2014).

## 3. PROGRAMS FOR ISING ANALYSIS OF FOLDING TRANSITIONS OF REPEAT PROTEINS

The overall workflow for our 1D-Ising analysis programs is shown in Figure 2A. Analysis is performed using three sequential python programs: (1) a program that converts data files to files containing numpy arrays, (2) a program that generates expressions for partition functions (equations 1-4) and fractional folding (equation 5) for each construct, and (3) a program that fits the converted data from the first program with a normalized expression based on fractional folding expressions generated from the second program. The third program also plots fitted data, performs bootstrap analysis, and analyzes parameter correlation. Each program is available on github (barricklab-at-jhu/Ising_programs) as a .py file that can be run directly in a terminal command line or in an IDE such as Anaconda (2016). In addition, the programs are combined in an interactive python notebook (.ipynb; Perez and Granger, 2007) that can be run in Jupyter (Randles et al., 2017).

**Figure 2.**
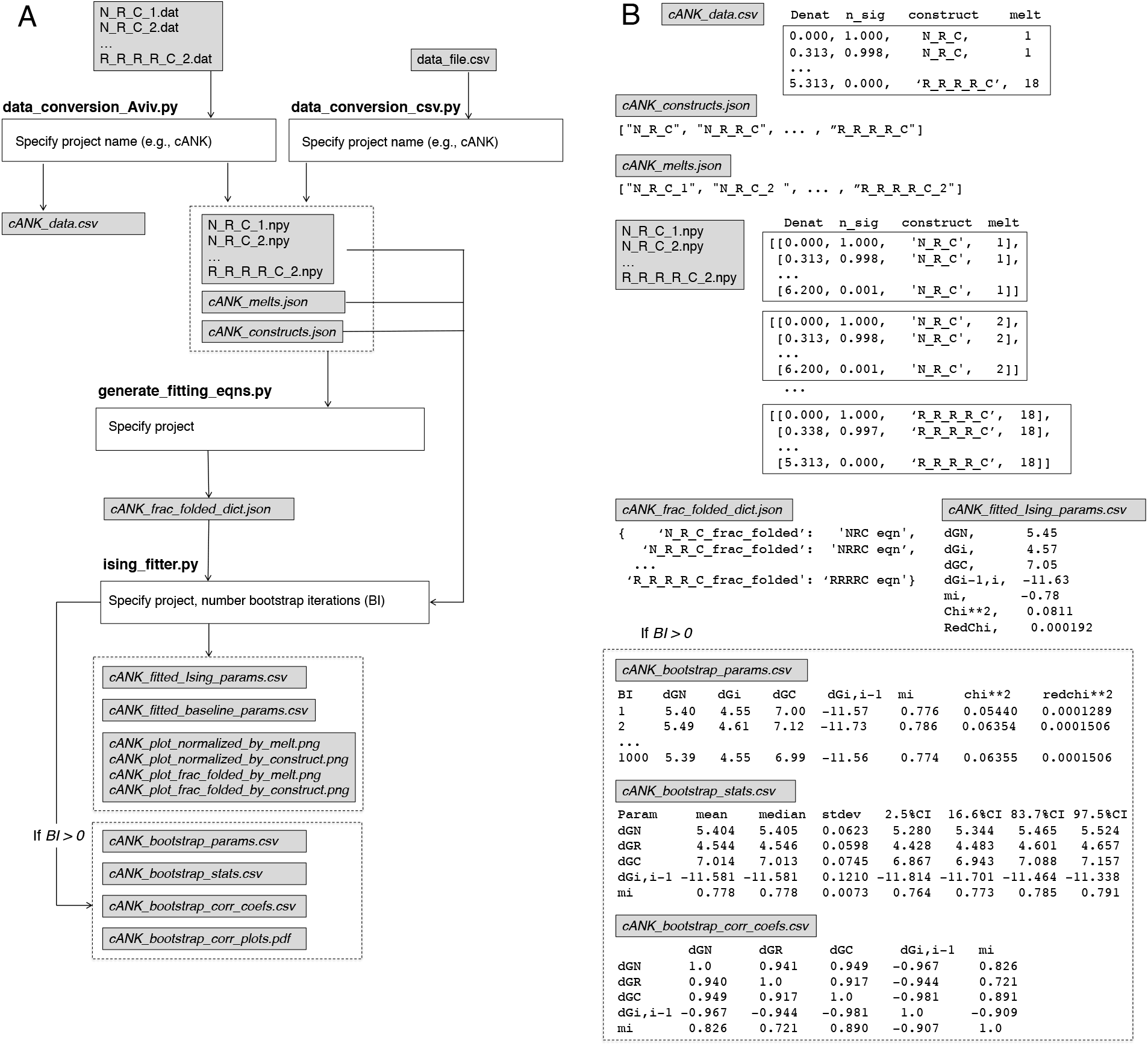
Flow-chart for 1D-Ising analysis of tandem-repeat unfolding transitions. Here, analysis of a capped homopolymeric series with three .py programs is illustrated. (A) Python programs are shown in white boxes along with required control parameters, input and output files are shown in grey boxes. (B) The format of input and output files. The first program (*data_preprocessing_Aviv.py*) converts data files to files containing numpy arrays and generates outputs lists of constructs and melts. The second program (*generate_fitting_equations.py*) uses the list of constructs to generate and output a dictionary of expressions for fraction folded (equation 5) for each construct. The third program (*ising_fitter.py*) uses these fraction-folded expressions, along with the melt and construct lists, to fit the expressions to the melt.npy data using nonlinear least squares. *ising_fitter.py* outputs fitted parameters to a csv file, as well as plots of raw and transformed data with fitted curves (see Figure 3). Following the fit, the user can run *BI* iterations of bootstrap analysis, which returns parameter values for each bootstrap iteration, statistics for bootstrap parameter distributions, a correlation matrix for bootstraped parameters, and plots of bootstrap histograms and parameter correlation (see Figure 4). Alternatively, all three programs can be run from a single combined Jupyter notebook.

### 3.1 Preprocessing data: data_preprocessing.py

The data preprocessing program generates a series of numpy arrays for each of the M melts and saves each to an *npy* file. In the process, this program writes out a list of constructs and a list of melts, which are used by the subsequent programs. To help identify output files, a “project” name should be specified in the data preprocessing program. This project name is used to name output files. It is convenient to use the name of the series to which the repeat constructs belong (such as “cANK” in the example in Figure 2).

There are two programs for data preprocessing that can be used for different types of input data. *data_preprocessing_csv.py* takes data in .csv format, which often requires some manual pre-processing, whereas *data_preprocessing_AVIV.py* takes input data in .dat format generated by Aviv spectrometers (the format of most of our denaturation data). In addition, *data_preprocessing_AVIV.py* outputs a single data csv file containing all melts, which can be combined with other csv data sets.

Input data files should be placed in the same folder as the jupyter notebook or the preprossessing program (and subsequent programs). For Aviv data files, each unfolding transition is in a separate file identified by the .dat extension. The root of each file name should give the structure of the repeat array for that melt, followed by an integer used to distinguish multiple melts of the same construct. For example, N_R_R_C_1.dat is the “first” melt for the construct NR_2_C (two internal R repeats flanked by N- and a C- capping repeats; Figure 1A). Likewise, R_X_R_C_2.dat is the “second” melt for the construct RXRC. For unfolding transitions in other formats, melts should be manually combined into a single .csv file containing four columns. The first column contains the denaturant concentrations for each melt, the second column contains spectroscopic values, the third column gives the construct in the format described above (e.g., N_R_R_C), and the fourth column gives an ID number identifying the data set, ranging from 1 to M (Figure 2B).

In the conversion process, data are normalized such that for each of the M melts the point with the largest signal intensity^4^ in each melt (*Y*_*max, k*_) is set to one, and the point with smallest intensity (*Y*_*min, k*_) is set to zero. The other data points in the *k*^*th*^ melt are scaled to these values:

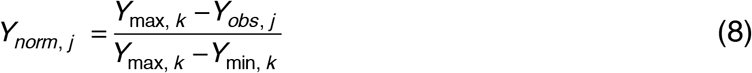

This normalization ensures that each melt has a similar influence in the fit (assuming each melt has the same number of points), which is important in cases where different melts have very different *Y* values. One potential pitfall is that if the starting *Y* values vary significantly from melt to melt, this scaling may produce significant differences in the absolute uncertainties in the normalized *Y* values of different melts. Such differences in uncertainties should be accounted for using weighting terms in the fitting program.

### 3.2. Generating fitting equations: generate_fitting_equations.py

Using the construct list generated by the data preprocessing program, the second program (*generate_fitting_equations.py*) builds a dictionary of partition functions (equation 4). From these partition functions, a dictionary of fraction folded expressions (equations 5 and 7) is built using the python *SymPy* module (which enables symbolic math manipulations, Meurer et al., 2017). A final and very important step in generating fraction folded expressions is the use of the *SymPy* command “simplify”, which significantly shortens the resulting expressions. For example, the simplify command reduces the expression for the fraction of folded NR_3_C from 467 to 227 characters. This reduction provides a significant speed-up during nonlinear least squares, which is especially important for bootstrap analysis (requiring hundreds to thousands of nonlinear-least squares optimizations). The dictionary of fraction folded expressions is output as a .json file for nonlinear least-squares fitting in the third program. Another feature built into *generate_fitting_equations.py* is a calculation of the rank of the coefficient matrix, which is useful evaluating whether the series of constructs being analyzed provides adequate constraints to fit the thermodynamic parameters (see Appendix 2). Results from the rank analysis are outputted to the screen, alerting the user to potential problems with the fit.

### 3.3. Fitting with an Ising model: ising_fitter.py

Using the converted data, construct list, and melt list generated by the data preprocessing program along with the fitting equations generated by the fitting equation program, the third program (*ising_fitter.py*) globally fits all of the folding transitions using nonlinear least-squares, by minimizing the sum of the squares of the residuals between normalized experimental *Y*_*norm, j*_ values from each of the M melts (equation 8) and the calculated values *Y*_*calc,k*_ (equations 5 and 7):

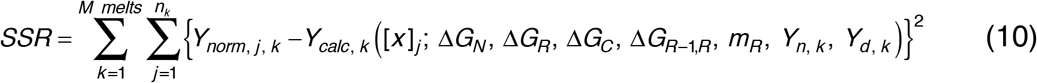

The terms in parentheses are the parameters to be optimized; in equation 10, they are those of model 1 (Table 1). The outer sum in equation 10 is over each of the M melts, and the inner sum is over each of the *n*_*k*_ denaturant concentrations in each melt. The fit is performed with the python *lmfit* module (Newville et al., 2019), which provides a number of useful fitting options including setting bounds for adjustable parameters.

The parameters that are optimized in equation 10 include global thermodynamic parameters, which apply to all melts and all constructs,^5^ and local baseline parameters, which apply individually to each melt (Table 1). For model 1, the thermodynamic parameters include intrinsic free energies for the N, internal R, and C repeats (Δ*G*_*N*_, Δ*G*_*R*_, and Δ*G*_*C*_), an interaction free energy between adjacent repeats (Δ*G*_*R−1,R*_), and a denaturant dependence for each repeat (*m*_*R*_, eq 5). The local parameters model a total of 2M baselines (a native and denatured baseline for each melt; *Y*_*n, k*_ and *Y*_*d, k*_ in equation 7). Each baseline is assumed to vary linearly with denaturant concentration; thus, each baseline requires two locally fitted parameters:

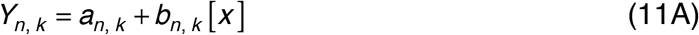

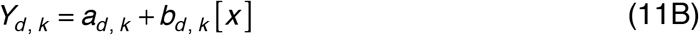

As is typical of nonlinear least-squares optimizations, initial guesses must be supplied to begin the search for optimal parameters. Finding satisfactory initial guesses for the thermodynamic parameters may require some care so that unfolding midpoints and slopes are reasonably approximated at the start of the search. Initial guesses for the baseline parameters are easily determined due to the normalization provided by the data preprocessing program: fitted values of *a*_*n, k*_ and *a*_*d, k*_ should be close to one and zero, respectively, and baseline slopes are often quite similar for all normalized melts. Thus, guesses for the 4*m* baseline parameters can usually be provided with just four parameters (*a*_*n*_, *b*_*n*_, *a*_*d*_, and *b*_*d*_).

From these initial guesses, *lmfit* uses the Marquardt-Levenberg algorithm to optimize the local and global parameters. The parameter set that minimizes the sum of the square of the residuals is then returned to the user as are plots of fitted data in various formats. Fitted thermodynamic and baseline parameters are also written to csv files, as are plots of data and fits (Figure 2).

### 3.4. Bootstrap analysis

To estimate uncertainties on fitted parameters and to explore parameter correlations, bootstrap analysis can be performed after the data have been fit. Bootstrap analysis is a resampling method in which sets of synthetic, or “bootstrapped”, data are generated with errors derived from the original data set (Efron, 1979; Johnson, 2008). Here we apply the bootstrap to the residuals of the fit, generating bootstrapped data sets by (1) using the best fitted parameters to estimate “true” values for each melt at each measured denaturant concentration (the *Y*_*calc, j, k*_ values in equation 10, using best-fitted parameters), (2) calculating residuals based on these *Y*_*calc, j, k*_ values (the expression in curly braces in equation 10), and (3) adding randomly selected residuals^6^ from step 2 to the true *Y*_*calc, j, k*_ values. Assuming the fitted Ising model provides a good description of the measured *Y*_*norm, j, k*_ values (specifically, that the *Y*_*calc, j, k*_ provides an good estimate of the error-free value of the measured *Y* value at denaturant concentration *j*, melt *k*, and that the errors in *Y*_*norm, j, k*_ are independent of *j* and *k*), these bootstrap data sets are an adequate approximation of independently determined data sets with the same structure as the observed data. Fitting each bootstrapped data set and comparing fitted parameter values provides an estimate of uncertainties for each fitted parameter.

One important parameter in bootstrap analysis is the number of bootstrap iterations performed. Ideally, thousands of iterations are performed, although for large data sets and complex models, performing thousands of bootstrap iterations may require more CPU time than is practical, especially on a desktop computer.^7^ The length of time required for bootstrap analysis can be significantly shortened by using parallel processing, either running on multiple cores within a laptop or desktop computer, or through shared processors within a cluster. We have created a version of the *ising_fitter.py* that runs bootstrapping in parallel using the python “*multiprocessing*” module (*ising_fitter_parallel.py,* also available on github). To help estimate the amount of time a given bootstrap analysis will take, the time required for the fit of the observed data is provided by *ising_fitter.py*, which is a good approximation of the time required for a single bootstrap iteration.

Upon completion of the fit of the data, *ising_fitter.py* prompts the user for the number of bootstrap iterations to perform. The fitter then performs the specified number of fits of bootstrap data, using the same initial guesses that were used for the initial fit. Thermodynamic parameters from each bootstrap iteration are written to a *bootstrap_params.csv* file, and a statistical summary of bootstrapped thermodynamic parameter values, including averages and measures of variation, are written to a *bootstrap_stats.csv* file (Figure 2).^8^

### 3.5. Correlation analysis

One advantage of the bootstrap method is that it permits visual inspection of parameter correlation. Analysis of parameter correlation reveals not only how much uncertainty is associated with fitted parameters, but the origins of that uncertainty. After the bootstrap analysis is completed, a grid of correlation plots is generated for all pairs of bootstrapped parameters along with a histogram of bootstrap values for each selected parameter. Linear Pearson correlation coefficients are calculated from each plot and are written to a *bootstrap_corr_coef.csv* file. Comparing the correlation plots for each parameter to the histogram of bootstrapped values emphasizes the important point that although parameter correlation increases parameter uncertainty, the uncertainty values reported from the bootstrap analysis include uncertainties resulting from correlation with all other parameters. Though a particular pair of parameters may show strong correlation, as long as the confidence intervals estimated from bootstrap analysis are tolerably low (i.e., the parameter histogram is suitably narrow), the correlation can be considered acceptable.

Rather, it is when fitted parameter values have intolerably large confidence intervals that correlation analysis is useful. In such cases, identifying the underlying correlations (in particular, pairs of parameters with large partial correlation coefficients; see below) that lead to large parameter uncertainties can be used to modify the model to avoid high correlation, or to include parameter constraints that are informed by independent information.

Although the correlation analysis described above reveals the extent to which pairs of parameters covary, these absolute correlations are sometimes indirect. Such indirect correlations result when each variable in a pair is each strongly correlated with a third. As a result, the two parameters in the pair show correlation with each other. To reveal the direct correlation among parameter pairs, “partial” correlation coefficients can be calculated in which correlation between a pair of parameters is calculated after correlation to all the other variable are accounted for. For a pair of variables X and Y in a data set with *n*+2 total variables, partial correlation coefficients are calculated by performing linear regressions separately for each variable X and Y with the *n* other variables **Z** = {Z_1_, Z_2_, …, Z_n_}, where {*X*, *Y*} ⊄ **Z**, determining the residuals for these two regressions (*r*_*X*,***Z***_ and *r*_*Y*,***Z***_), and determining the correlation coefficients for the residuals *r*_*X*,***Z***_ and *r*_*Y*,***Z***_ (referred to as *ρ*_*X,Y•Z*_):

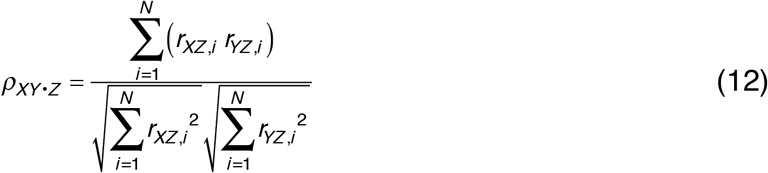

Using the values of bootstrapped thermodynamic Ising parameters in *boostrap_params.csv*, partial correlation coefficients are calculated using the program “*partial_correlation.py*”. This program uses the *Pingouin* statistical package (Vallat, 2018) to calculate partial correlation coefficients, and outputs them to the file *bootstrap_partial_corr.csv*.

## 4. A FIVE-PARAMETER (MODEL 1) FIT OF A CAPPED HOMOPOLYMER SERIES

Using the fitting programs above, we have performed Ising fitting and bootstrap analysis on unfolding transitions of a series of capped homopolymeric consensus ankyrin repeat constructs described previously (Aksel and Barrick, 2009). The constructs in this series match those in Figure 1A (model 1, Table 1), ranging in length from three to five repeats, and containing either N-terminal caps, C-terminal caps, or both. For each construct, there are two melts (technical replicates), such that each construct has a similar impact on the fit. Likewise, for each capping pattern (NR_x_, R_x_C, NR_x_C), there are three constructs of different length (x=3, 4, 5), so that the two caps have a similar impact on the fit. As a whole, the data set comprises 18 melts with a total of 499 observations. There are five fitted global thermodynamic parameters (ΔG_N_, ΔG_R_, ΔG_C_, ΔG_R−1,R_, and m_R_), and 72 local baseline parameters (four for each melt). Thus, there are 499–5–72 = 422 degrees of freedom (DOF). The fit took about 5 seconds of wall time on a MacBook Pro using a single 2.3 GHz Intel Core i9 processor (although this time depends on values of the initial guesses).

Overall, the fit of model 1 to the data is quite good. This can be seen in Figure 3A and B, where the normalized data and fitted curves are in good agreement. Inspection of normalized curves is important because this is the space in which the fit is performed. Highly sloped baselines, which are often a result of inadequate baseline sampling (and which often increase uncertainties on fitted thermodynamic parameters), are apparent upon inspection of normalized fits. In contrast, baseline slopes are hidden in fraction folded curves (Figure 3C and D). For melts in which fitted baseline slopes are unreasonably high, limits can be imposed on baseline slopes using the *lmfit* module by providing *min* and *max* values during initial parameter assignments.^9^ The quality of the fit in Figure 3 is also indicated by the reduced sum of squares of the residuals (RSSR = SSR/DOF = 0.00019 in Figure 3A). Since the normalized unfolding transitions span one unit of signal change, this RSSR value reflects an average squared residual of about 0.019 percent, or an average residual of about 1.4 percent. This residual is within the error resulting from the circular dichroism measurements from which the normalized signal is derived.

**Figure 3.**
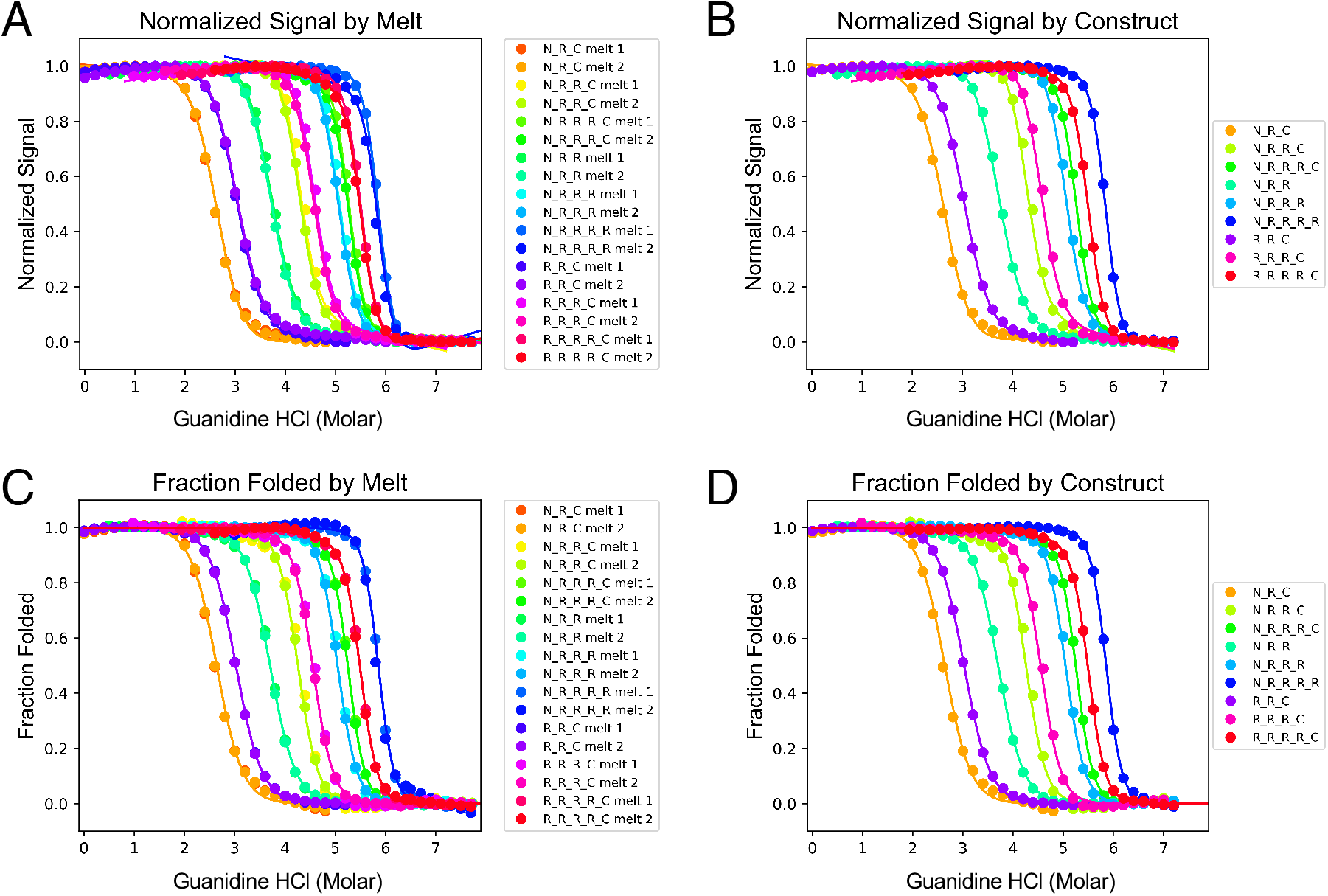
Fitted unfolding transitions for capped homopolymeric ankyrin arrays. Data are from (Aksel et al., 2011) and are fitted with model 1 (Table 1). Plots are direct outputs from the plotting section of the fitter. Data and fitted curves are plotted both as normalized signal (A and B, the space in which least-squares minimization is carried out) and as fraction folded (C and D). In the fraction folded representation, fitted curves for different melts of the same construct are identical (because a single set of thermodynamic parameters is fitted globally), but the data are not (due to random errors). In contrast, fitted curves to normalized data differ, since each curve has its own baseline parameters. Plots (A) and (C) show all fitted melts; plots (B) and (D) show one representative melt from each construct.

To estimate uncertainties on the fitted thermodynamic parameters, 1000 bootstrap iterations were performed. In absolute terms, uncertainties on the thermodynamic parameters are quite small for the fit in Figure 3: 95 percent confidence intervals^10^ of bootstrapped free energies are about 0.2 kcal mol^−1^ (Table 2). Bootstrap estimates for all thermodynamic parameters have unimodal and roughly normal distributions (diagonal histograms, Figure 4), consistent with a single well-defined minimum in SSR over the fitted parameter space. For each parameter, uncertainties can be estimated from the standard deviation of bootstrap values or from locating the parameter values that separate the low and high tails of the bootstrap distribution at 67 or 95 percent (Table 2). An advantage of the latter approach is that no assumptions made on the symmetry of the error. For the five thermodynamic parameters in model 1, the 67 and 95 percent confidence intervals are within 0.1 percent of the mean plus or minus one or two standard deviations, as is expected for normally distributed parameter estimates. Large differences between the standard deviation and confidence intervals may reflect fitting pathologies such as one-sided parameter bounds, and multiple least-squares minima in parameter space (Johnson, 1983), which should be apparent in bootstrap parameter histograms. Identifying such pathologies is important, since it almost certainly indicates that the model is not adequately constrained by the constructs being fitted, and that either the model or the data set should be modified.

**Table 2.** Fitted global thermodynamic parameters and bootstrap statistics.

|  |  | Bootstrap statistics <sup>b</sup> |  |  |  |  |  |
| --- | --- | --- | --- | --- | --- | --- | --- |
| Parameter<sup>a</sup> | Best fit value | Mean | Standard deviation | lower 95% CI <sup>c</sup> | lower 67% CI <sup>c</sup> | upper 67% CI <sup>c</sup> | upper 95% CI <sup>c</sup> |
| *Model 1* |  |  |  |  |  |  |  |
| $\Delta G_N$ | 5.47 | 5.41 | 0.062 | 5.28 | 5.34 | 5.46 | 5.52 |
| $\Delta G_R$ | 4.57 | 4.54 | 0.060 | 4.43 | 4.48 | 4.60 | 4.66 |
| $\Delta G_C$ | 7.06 | 7.01 | 0.075 | 6.87 | 6.94 | 7.09 | 7.16 |
| $\Delta G_{R-1,R}$ | -11.63 | -11.58 | 0.12 | -11.81 | -11.70 | -11.46 | -11.34 |
| $m_R$ | -0.78 | -0.78 | 0.0073 | -0.764 | -0.773 | -0.785 | -0.791 |
| *Model 2* |  |  |  |  |  |  |  |
| $\Delta G_N$ | 4.89 | 4.89 | 0.064 | 4.77 | 4.82 | 4.95 | 5.01 |
| $\Delta G_R$ | 4.08 | 4.08 | 0.065 | 3.96 | 4.02 | 4.14 | 4.22 |
| $\Delta G_X$ | 5.64 | 5.64 | 0.091 | 5.46 | 5.55 | 5.73 | 5.82 |
| $\Delta G_C$ | 6.34 | 6.34 | 0.073 | 6.20 | 6.27 | 6.41 | 6.49 |
| $\Delta G_{R-1,R}$ | -10.46 | -10.46 | 0.12 | -10.71 | -10.58 | -10.36 | -10.25 |
| $\Delta G_{X-1,X}$ | -10.16 | -10.17 | 0.12 | -10.39 | -10.29 | -10.05 | -9.94 |
| $\Delta G_{R-1,X}$ | -9.42 | -9.42 | 0.11 | -9.64 | -9.52 | -9.33 | -9.22 |
| $\Delta G_{X-1,R}$ | -10.82 | -10.83 | 0.12 | -11.07 | -10.95 | -10.72 | -10.59 |
| $m_R$ | 0.71 | 0.708 | 0.0071 | 0.69 | 0.70 | 0.71 | 0.72 |
| $m_X$ | 0.97 | 0.98 | 0.021 | 0.93 | 0.95 | 0.10 | 1.02 |

<sup>a</sup> $\Delta G$ values in kcal mol<sup>-1</sup>; $m$ -values in kcal mol<sup>-1</sup> M denaturant<sup>-1</sup>. <sup>b</sup> Values from 1000 bootstrap iterations. <sup>c</sup> CI, confidence intervals.

**Figure 4.**
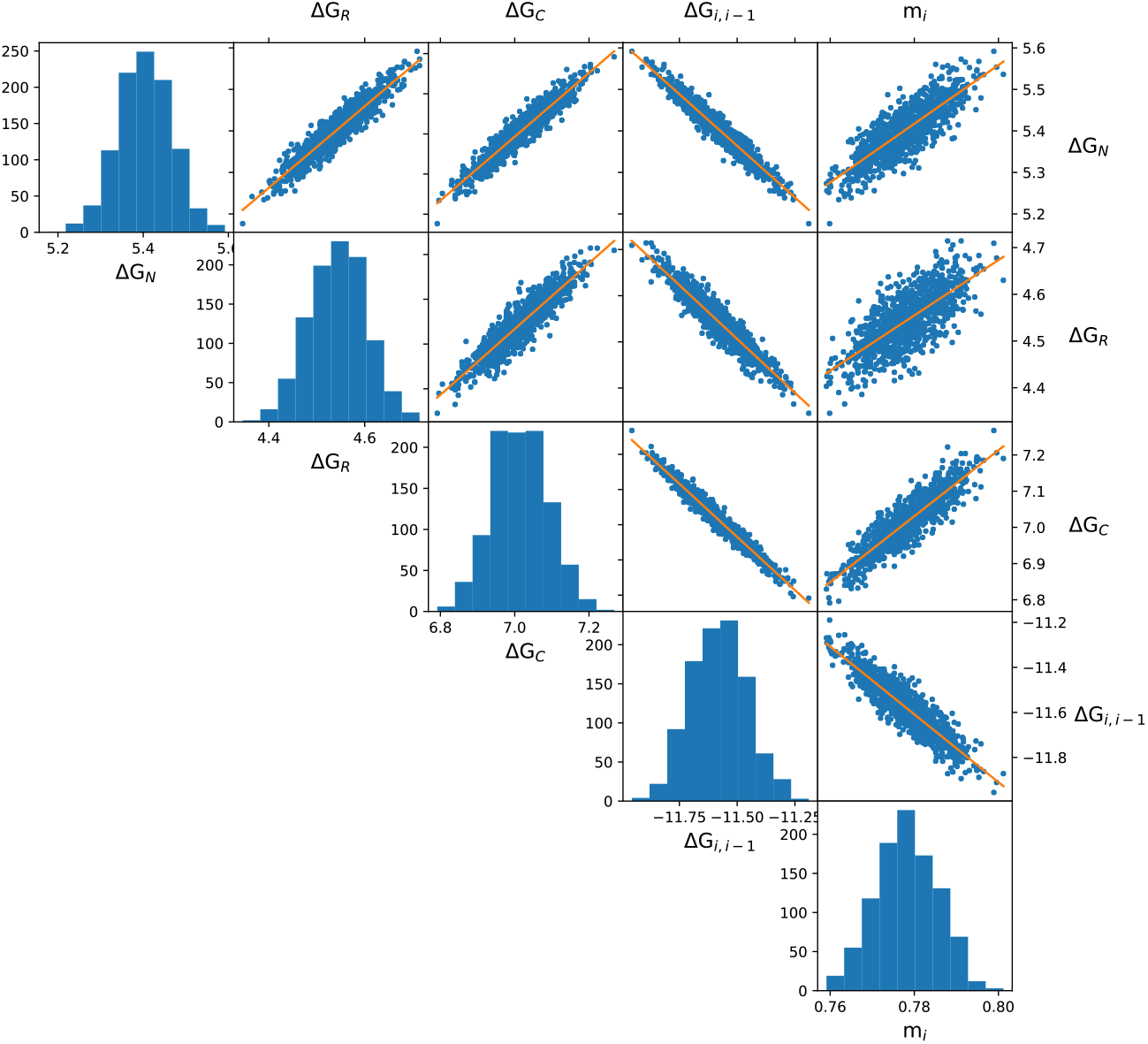
Bootstrap histograms and correlation plots of fitted thermodynamic parameters for capped homopolymeric ankyrin arrays. Capped consensus ankyrin homopolymer data from Figure 3 was used along with best-fitted parameters from Model 1 to generate 1000 bootstrap fits. The lower diagonal panels show histograms for bootstrapped thermodynamic parameters. The upper triangular panels show correlations among pairs of thermodynamic parameters, along with a best-fitted line to bootstrapped pairs.

As described above, parameter correlation can be a significant source of uncertainties in fitted parameters. Even for the simplest types of fits, strong correlations are common, such as the fitted slope and intercept parameters in linear regression against a single independent variable (Johnson, 1983), and the ΔG° and m-values describing simple two-state folding (see Appendix 1). The bootstrap analysis allows correlations between pairs of parameters to be clearly visualized in a scatter plot in which each point represents a pair of parameter values from the same bootstrap iteration (Figure 4). Correlations between bootstrap parameter pairs can also be seen in the correlation coefficient matrix that is generated by *ising_fitter.py* (as *bootstrap_corr_coef.csv*, Table 3).

**Table 3.** Bootstrap parameter correlation coefficients for Model 1^*a*^.

| Parameters | $\Delta G_N$ | $\Delta G_R$ | $\Delta G_C$ | $\Delta G_{R-1,R}$ | $m_i$ |
| --- | --- | --- | --- | --- | --- |
| $\Delta G_N$ | 1 | 0.941 | 0.949 | -0.967 | 0.826 |
| $\Delta G_R$ | -0.710 | 1 | 0.917 | -0.947 | 0.721 |
| $\Delta G_C$ | -0.485 | -0.699 | 1 | -0.981 | 0.891 |
| $\Delta G_{R-1,R}$ | -0.782 | -0.986 | -0.766 | 1 | -0.909 |
| $m_R$ | -0.753 | -0.988 | -0.689 | -0.989 | 1 |

<sup>a</sup> Values from 1000 bootstrap iterations. Values above the diagonal are Pearson correlation coefficients between pairs of variables. Values below the diagonal (in red) are Pearson partial correlation coefficients between pairs of variables, where the effects of the other variables in the table are factored out (see text).

For the capped homopolymer fit (Figure 3), the strongest correlations are between the interfacial free energy (ΔG_R−1,R_) and the three intrinsic free energies (ΔG_N_, ΔG_R_, and ΔG_C_; Table 3); these correlations are negative (Figure 4).

This negative correlation between intrinsic and interfacial free energies is expected—an increase in ΔG_R−1,R_ (i.e., a decrease in the interfacial free energy), which would decrease midpoints of unfolding transitions, can be compensated by decreasing ΔG_N_, ΔG_C_, and especially ΔG_R_ (since this term contributes three times as much, on average, to the stabilities of the constructs in Figure 1 as ΔG_N_ and ΔG_C_).

In addition, there appear to be significant positive correlations among ΔG_N_, ΔG_R_, and ΔG_C_ (Table 3, Figure 4). These correlations are unexpected—if one intrinsic free energy parameter increases, the others would be expected to decrease to maintain the midpoint of the folding transition. Thus, it seems likely that these apparent correlations are indirect—since each intrinsic free energy is negatively correlated with ΔG_R−1,R_, each may end up positively correlated with the other two. To test for indirect correlation, we calculated partial correlation coefficients among the thermodynamic bootstrap parameter values, which reveal the correlation between two parameters when correlations to the other parameters are factored out (Table 3, lower triangle). Based on partial coefficients, the negative correlation between ΔG_R−1,R_ and ΔG_R_ remains high (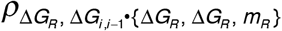 is actually more negative than the total correlation coefficient between these two parameters; Table 3), but the correlations between ΔG_R−1,R_ and ΔG_N_ and ΔG_C_ are decreased. Moreover, the correlations among the intrinsic free energies (ΔG_N_, ΔG_R_, and ΔG_C_) are also decreased, and the signs of the partial correlation coefficients are negative, matching expectations.

Partial correlation analysis reveals a similar indirect correlation between the bootstrapped m_R_-values and the three bootstrapped intrinsic free energy parameters. Though the total correlation coefficients of all four free energy bootstrapped parameters with m_R_ are large (Figure 4), only ΔG_R−1,R_ shows the expected negative correlation (negative correlation is expected since an increase in m_R_ should increase the stability in the absence of denaturant, decreasing ΔG values; see Appendix I). The three intrinsic free energy parameters show unexpected positive total correlations with m_R_, with slightly larger correlation coefficients for ΔG_N_ and ΔG_C_ than for ΔG_R_. The partial correlation coefficients for these three parameter pairs 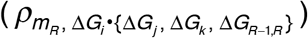 have reversed sign, with the strongest negative correlation between m_R_ and ΔG_R_, suggesting the positive total correlation is indirect. In contrast, the correlation between m_R_ and ΔG_R,R−1_ remains large and negative.

Overall, the picture that emerges is that there is a strong negative direct correlation between ΔG_R_ and ΔG_R−1,R_, and also between those two parameters and m_R_. ΔG_N_ and ΔG_C_ have weaker direct negative correlations to these three parameters, and weaker correlations to each other. Again, it is worth pointing out that despite these correlations, ΔG_R_ and ΔG_R−1,R_ are determined within 0.2 kcal mol^−1^. As we have previously described, fitted values of ΔG_R_ and ΔG_R−1,R_ are consistent with a strong positive cooperativity in which intrinsically unstable repeats are driven to fold by strongly stabilizing interfaces with folded neighbor repeats (Aksel and Barrick, 2014; Aksel et al., 2011).

## 6. A TEN-PARAMETER (MODEL 2) FIT OF A CAPPED HETEROPOLYMER SERIES

To illustrate how the Ising fitting programs perform with a more complex heteropolymeric model and to examine the effects of a mutation on intrinsic and interfacial stabilities, we performed Ising fitting and bootstrap analysis on folding transitions of a series of capped consensus ankyrin repeats in which a conserved threonine from one or two R repeats is substituted with a valine at position 7 in the ankyrin TPLH motif (referred to here as T7V). The T7V substitution has previously been shown to destabilize a four-repeat NXRC construct (where repeat X is an R-type repeat that harbors the point substitution) by 2.6 kcal mol^−1^ using a simple two-state model (Preimesberger et al., 2015). By combining the X-substitution series (Figure 1B) with the homopolymeric capped constructs analyzed with model 1 above,^11^ we were able to fit a model for heteropolymeric capped repeats containing ten thermodynamic parameters (model 2).

To accurately determine the intrinsic and interfacial free energies, it is essential that the constructs being analyzed provide adequate constraints on these quantities during the fit. For capped homopolymer series, having the sufficient constructs to constrain ΔG_N_, ΔG_R_, ΔG_C_, and ΔG_R−1,R_ is fairly intuitive^12^, but for capped heteropolymeric series, the complexity of the model and the number of thermodynamic parameters can obscure this sufficiency. To check whether or not the intrinsic and interfacial free energy parameters of the model are adequately determined by the constructs in the data set, the rank of the coefficient matrix defined by the series of constructs is evaluated by *generate_fitting_equations.py*. If the coefficient matrix has full column rank, the constructs adequately define the thermodynamic parameters (Appendix 2). Indeed, the combined constructs from Figure 1A and B pass the rank test for the parameters in model 2 (Table 1), permitting us to determine four intrinsic free energies (ΔG_N_, ΔG_R_, ΔG_X_, and ΔG_C_) and four interfacial free energies (ΔG_R−1,R_, ΔG_X−1,X_,ΔG_R−1,X_, and ΔG_X−1,R_). As a whole, the data set comprises 36 melts with a total of 1057 observations. In addition to the eight global thermodynamic parameters above, there are two m-values that give the denaturant dependences of ΔG_R_ and ΔG_X_ (m_R_ and m_X_; equation 6), and 144 baseline parameters. Thus, there are 1057–10–144 = 903 degrees of freedom. The fit took about 10 seconds of wall-time, about twice that for model 1, which involves half the number of melts.

Overall, the heteropolymeric capped repeat data set is reasonably well-fitted by model 2 (Figure S1). Fitted values for thermodynamic parameters shared between models 1 and 2 are within about 0.5 kcal mol^−1^ for intrinsic folding energies and 1 kcal mol^−1^ for the interfacial energy. The reduced sum of squares of the residuals is 0.00038, corresponding to an average residual of about 2 percent per point. Uncertainties in fitted thermodynamic parameters for the model 2 fit are quite similar to those from model 1, both for parameters that are shared by the two models and for the parameters that are unique to model 2 (Table 2). An exception is m_X_, which has about three times the uncertainty as m_R_. This is not unexpected since there are significantly fewer repeats in the fitted data set that have an m_X_-denaturant dependence (the X repeats) than repeats that have an m_R_ dependences (N, R, and C repeats).

Although model 2 contains more fitted parameters than model 1, correlations among fitted bootstrap parameters seem to be somewhat reduced. There are modest decreases in both total and partial correlation coefficients for most of the parameter pairs that are shared by both models (compare Tables S1 and 3). This difference is borne out visually in a comparison of the two sets of correlation plots (Figures S2, 4). The weakest correlations involve the two parameters in model 2 that are least well determined: m_x_ and in particular, ΔG_X_. This relationship between parameter correlation and uncertainty underscores the point made above that strong parameter correlation is not a direct proxy for high parameter uncertainty. Rather, at least in the examples shown here, high parameter uncertainty appears to mask correlation, whereas low parameter uncertainty reveals correlation.

The additional parameters in model 2 provide a quantitative measure of how the point-substitution affects intrinsic and interfacial stability. Comparing ΔG_X_ with ΔG_R_ reveals that intrinsic stability is decreased by 1.6 kcal mol^−1^. This is consistent with the loss of an internal hydrogen bond between the threonine (O_γ_)H and the *i+3* histidine side-chain N_δ_1 (Preimesberger et al., 2015). This hydrogen bond is part of a hydrogen bond network that includes a second, bifurcated hydrogen bond from the *i+3* histidine N_δ_1 to the threonine main-chain (N)H, and from the *i+3* histidine N_ε_2 (N)H to the main-chain C=O from the residue preceding the threonine in the next repeat (asparagine *i+32*; Figure 5A).

**Figure 5.**
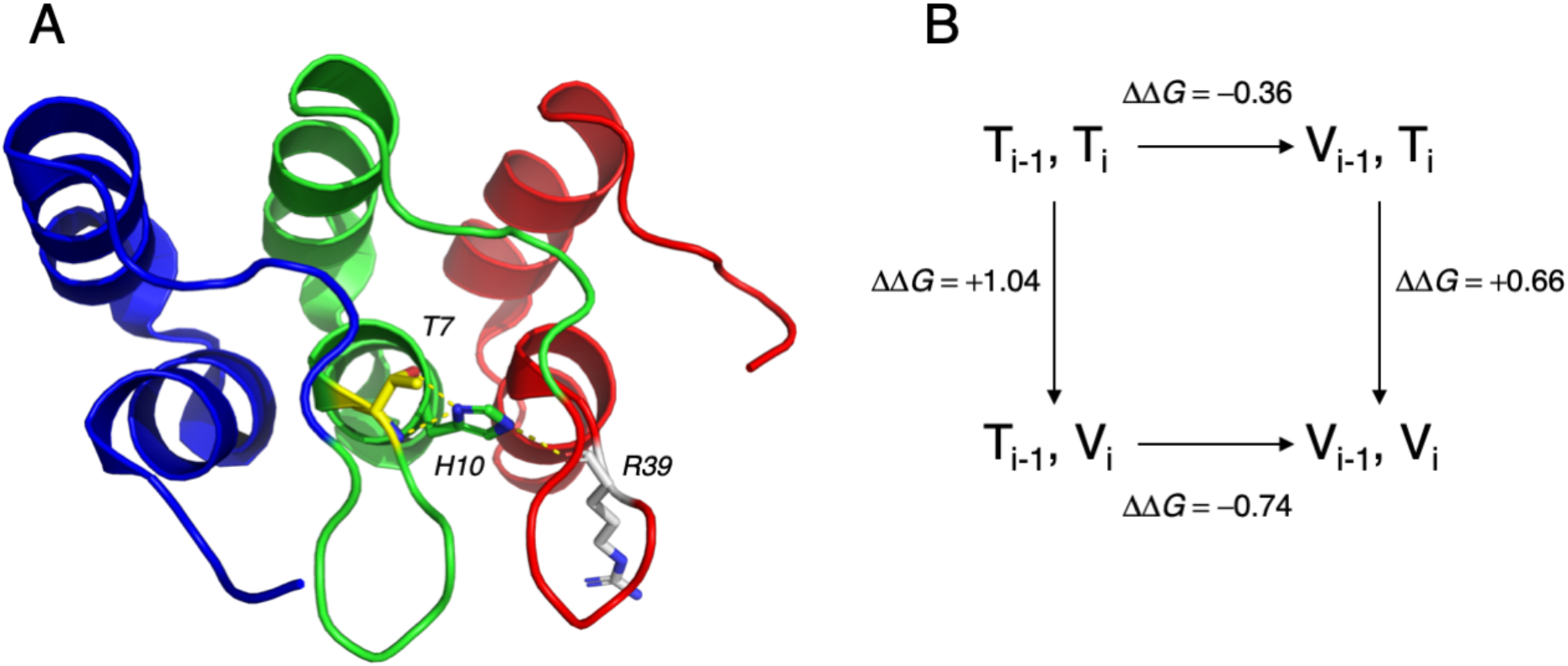
Threonine 7, histidine 10 and their hydrogen bonding patterns within and between ankyrin repeats. (A) Hydrogen bonding pattern in a consensus ankyrin array from Mosavi et al. (1N0R.pdb, Mosavi et al., 2002); note that the histidine ring has been flipped by 180° around χ_2_ to match the bonding pattern determined by Preimesberger et al., 2015). Threonine at position 7 (yellow) forms a bifurcated hydrogen bond to the Nδ_1_ of histidine 10 via its main-chain NH and side-chain OH within the same repeat. In turn, histidine 10 hydrogen bonds to a main-chain CO in the C-terminal neighboring repeat. (B) Thermodynamic cycle involving changes to interfacial interaction energies (Δ*G*_*i−1,i*_) in response to T7V substitution in adjacent repeats. For each arrow, the ΔΔG value is the difference between the interaction energies in the substituted (arrowhead) and unsubstituted interfaces (arrowtail). For example, for the top arrow, ΔΔ*G* = Δ*G*_*X−1,R*_ − Δ*G*_*R−1,R*_. The difference between ΔΔG values on the top and bottom (and equivalently, left and right) arrows gives the interaction energy between the two substitutions, here −0.4 kcal mol^−1^.

Comparing ΔG_X−1,X_ with ΔG_R−1,R_ provides one measure of how interfacial stability is affected by point substitution. This comparison indicates that the interface may be weakly destabilized by valine substitution in adjacent repeats by 0.3 kcal mol^−1^, although this value is only slightly beyond the 95% confidence intervals (0.12 kcal mol^−1^ for each free energy). However, by combining substitutions to R-repeats (X) with unsubstituted N, R, and C repeats as in Figure 1B, we can also determine interfacial free energies between an N-terminal R repeat and an adjacent X repeat (ΔG_R−1,X_), and an N-terminal X repeat and an adjacent R repeat (ΔG_X−1,R_). In other words, we can compare the energetic effects of single point substitution in the N- (*i, i−1*) and C-terminal (*i, i+1*) directions. Comparison of ΔG_R−1,X_ and ΔG_X−1,R_ with ΔG_R−1,R_ shows that the N-terminal interface is destabilized by valine substitution by +1.0 kcal mol^−1^, but that this stabilization is partly compensated by a stabilization of the C-terminal interface by −0.4 kcal mol^−1^ (Figure 5A). The modest stabilization of the C-terminal interface may reflect a re-orientation of the *i+3* histidine side-chain in the substituted repeat to optimize the two remaining hydrogen bonds (Preimesberger et al., 2015).

Finally, the fact that the two interfacial free energy perturbations do not sum to the perturbation resulting from a tandem substitution identifies a thermodynamic coupling between adjacent repeats. This coupling can be represented in a thermodynamic cycle (Figure 5B). Although the magnitude of the coupling is moderate (~0.4 kcal mol^−1^), the sign of the coupling favors interfaces with the same residue at repeat position 7 of adjacent repeats. Pairs of threonine residues at repeat position seven result in the most stable interface; although pairs of valines at position seven result in a destabilized interface, a valine pair is more stable than would be expected based on the interfacial energy perturbations from single-repeat substitutions. In other words, pairs of valines at position seven are mutually stabilizing. This type of long-range sequence coupling should serve to reinforce the repetitive nature of ankyrin sequence motif. The approach described here provides a means to identify other positions in the 33-residue ankyrin motif that are similarly coupled.

## 7. SUMMARY

Understanding cooperativity in protein folding, which is determined by the distribution of local and long-range stabilities, is likely to have important functional consequences in terms of avoiding misfolding and in allosteric functional transitions. The analysis of tandem repeat protein arrays with a 1D Ising model provides a unique approach to quantifying cooperativity in terms of local stabilities and long-range coupling free energies. The collection of programs presented here, which are available on github, provide a robust platform for data processing, fitting, plotting, and analysis of parameter uncertainties and correlations. Given the complexity of the models, careful inspection of fitted parameter uncertainties is an essential part of the analysis. Models can easily be adapted to evaluate the effects of point-substitutions in intrinsic and interfacial energies, providing insights into the structural basis of cooperativity.

## Acknowledgments

KG, JM, SK, and MP were supported by NIH training grant 2T32 GM008403. KS was supported by NIH training grant 5T32 GM007231. This work was supported by NIH grant R01 GM068462 to DB. The authors declare no conflict of interest.

## Appendix I. Correlation between fitted parameters in simple two-state unfolding analysis

The Ising analysis of repeat protein folding transitions above shows that there are significant correlations between fitted parameters, based on bootstrap analysis. For example, in the homopolymer fit using model 1, there are strong direct correlations between ΔG_R_ and ΔG_R−1,R_ (Table 3). Here we will use the Ising programs outlined in Figure 1 along with melts from a single construct to demonstrate that this type of correlation is not unique to multiparameter Ising fits, but is also endemic (but perhaps often overlooked) in fitting of simple two-state unfolding transitions.

To directly investigate correlation using a two-state model, the Ising fitting procedure outlined in Figure 1 was performed using two melts from just one construct, NRC. To fit with a two-state model, the Aviv .dat files were renamed R_1.dat and R_2.dat. As a result, *generate_fitting_equations.py* builds a single partition function of the form

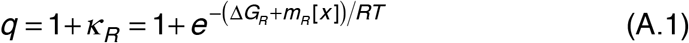

Differentiating with respect to *κ*_*R*_ and converting the result to an expression for *Y*_*calc*_ (equations 5 and 7) gives a simple two-state model that can be fitted to the renamed NRC melts and subjected to two-parameter (ΔG_R_ and m_R_) bootstrap analysis (Figure A.1).

**Figure A.1.**
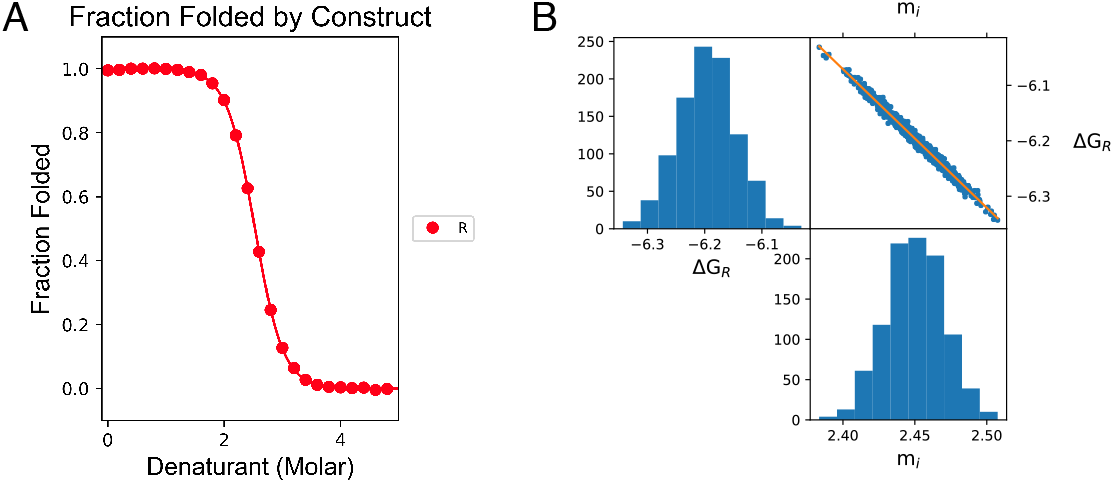
Parameter correlation in a simple two-state fit. NRC (Figure 1A) was fitted with a two-state model (deriving from equation A.1) using the Ising programs described here. (A) The NRC folding transition is well-fitted by the two-state model. (B) Bootstrap analysis (1000 iterations) shows that the two fitted thermodynamic parameters are determined with narrow confidence intervals, but they are tightly correlated.

The fits are excellent (Figure A.1A), and bootstrapped ΔG_R_ and m_R_-values are determined with narrow confidence intervals (within 1 to 2 percent). However, these bootstrapped parameters are very strongly correlated (Figure A.1B): the correlation coefficient between ΔG_R_ and m_R_ is −0.993. Note that because there are just two thermodynamic parameters in the correlation analysis, partial and total correlation coefficients are equivalent.

## Appendix II. Rank analysis of the thermodynamic parameters in model 2

To accurately determine intrinsic and interfacial free energies from Ising analysis of repeat protein folding, the constructs being analyzed must present enough structural variation to accurately determine the model’s parameters. For example, model 2 has eight free energies (four intrinsic folding energies and four interfacial energies). To accurately determine these parameters, the set of constructs analyzed (the combined constructs in Figure 1A and B for the analysis described here) must provide uncorrelated variations in structural features that determine each parameter. For complex models and large data sets, it is not always obvious that a given data set is adequate to determine the parameters in a given model.

Here we present a simple method to check whether a data set provides enough structural variation to adequately constrain the parameters for a specific model. Since the folding free energy for each construct is expressed as a sum of Ising parameters, the compatibility of a thermodynamic model can be tested by evaluating the set of linear free energy equations defined by the model for each of the constructs being analyzed (Aksel and Barrick, 2009). The constructs analyzed using model 2 define 15 linear free energy equations (Figure A.2). If the system of equations has an empty null space, then the system of equations has a unique solution. An easy way to test whether the null space of a system of equations is empty is to compute the matrix rank. If the rank of a coefficient matrix is equal to the number of columns in the matrix, the null space is empty and a unique solution to the system of equations exists. The *generate_fitting_equations* program performs this rank analysis, and reports the rank (and its sufficiency/insufficiency) during run time. In Figure A.2, which gives the free energy equations for the constructs used for Ising analysis with model 2 in matrix form, the coefficient matrix has a full column rank of 8. Thus, this set of constructs adequately defines the unknown parameters in model 2.

**Figure A.2.**
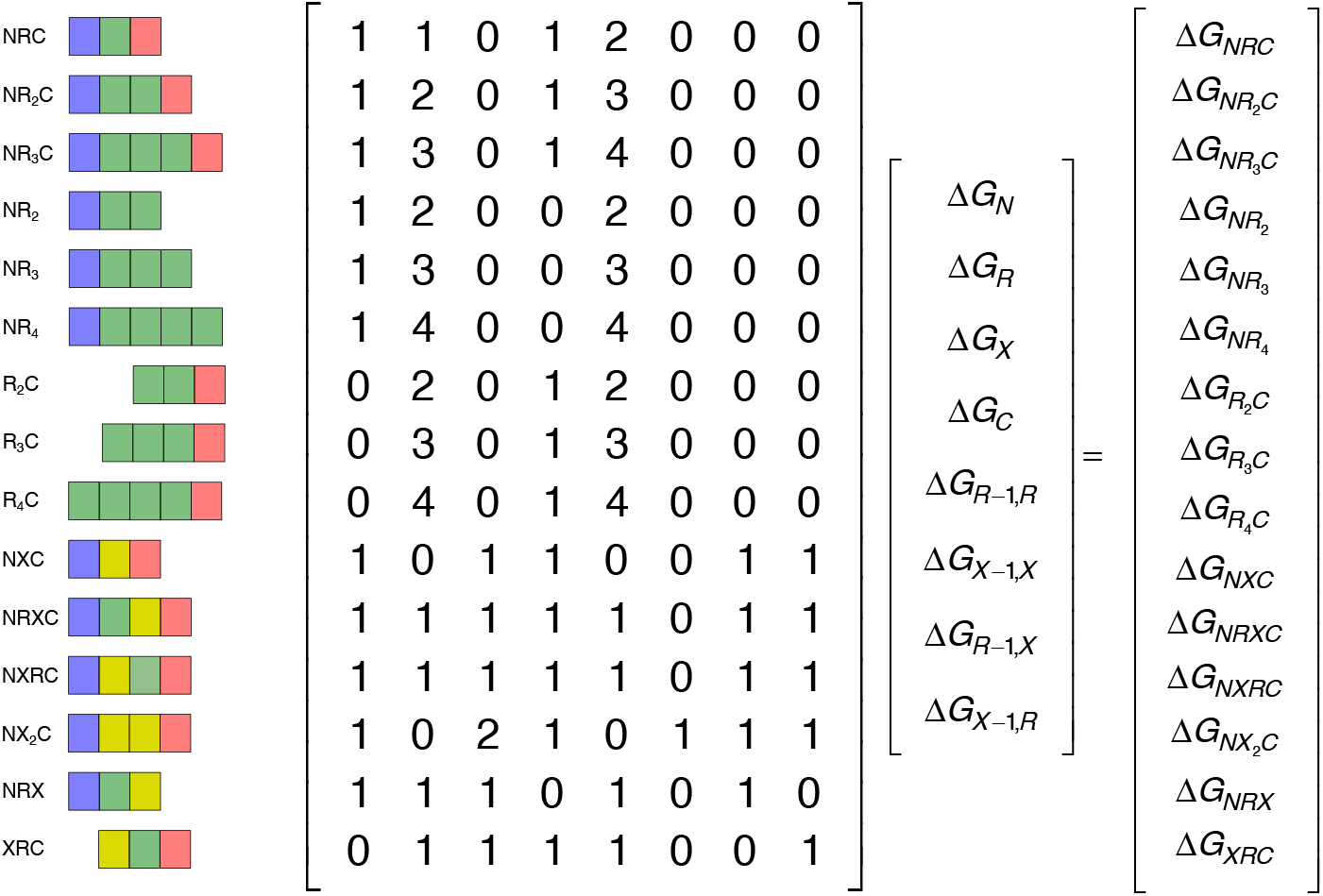
A matrix representation of the equations for repeat protein stability with the Ising model. The constructs that are used to fit the parameters from model 2 are shown on the left. These constructs define a set of 15 linear equations in which the free energies of the fully folded state relative to the unfolded state (the vector on the right-hand side) can be represented as a sum of intrinsic and interfacial free energies (the 8 by 1 vector on the left-hand side), weighted by the number of times each term is represented in a given construct (the 8 by 15 coefficient matrix). To have a unique solution, each column in the coefficient matrix must be independent of the others, that is, the matrix must have full column rank (eight in the example here).

In contrast, a model that treats interfaces between X-repeats and capping repeats as unique (with terms ΔG_NX_ and ΔG_XC_) cannot be uniquely fitted by the data set in Figure A.2. The corresponding coefficient matrix has ten columns, but the rank remains eight. Although the Ising fitter converges on a least-squares solution for the ten-repeat model, the fitted parameter values differ significantly from values from simpler models, and the confidence intervals on many of the parameters are extremely broad. Such cases require either simplification of the model (to model 2, for example), or in favorable cases, inclusion of additional constructs that provide the missing structural variation.

## Footnotes

1 As with ghosts, there is no such thing as a zeroth repeat.

2 In principle, denaturant dependence of the interfaces could be included by introducing an interfacial m-value through an equation analogous to (6), although in practice, *m*_*i*_ and *m*_*i−1,i*_ are strongly coupled.

3 In subsequent expressions, it will be assumed that the kth melt belongs to the set of M total melts, e.g., *Y*_*calc, k∈M*_ → *Y*_*calc, k*_.

4 For CD measurements in the far-UV, this is typically the largest *negative* value.

5 An exception is the intrinsic free energies of folding of the capping repeats (Δ*G*_*N*_ and Δ*G*_*C*_), which apply to all repeats that contain N- and C-terminal caps.

6 Random selection with replacement.

7 Factors that increase the amount of bootstrap CPU time are the number of melts, which determines the number of fitted baseline parameters), the number of thermodynamic parameters (e.g., compare Mode 2 with Model 1, Table 1), and the extent of parameter correlation.

8 Although all fitted parameters (thermodynamic and baseline parameters) are optimized in each bootstrap iteration, we are typically interested in uncertainties and correlations among the thermodynamic parameters. If the uncertainties in fitted baseline parameters are desired, the bootstrap portion of *ising_fitter.py* can easily be modified to include these values for statistical analysis.

9 In most cases, a preferable remedy for a poor baseline is to collect a melt with a better baseline. However, this is not always possible for marginally or highly stable constructs.

10 Given that the bootstrap approach misses some sources of error, such as systematic variations among different data sets, 95% confidence intervals seem a safer and more appropriate estimate of uncertainties in fitted parameters than the more common 67% limits.

11 For the combined data set, we included three melts for each of the six heteropolymer constructs in Figure 1B, and two melts for each of the nine homopolymer constructs in Figure 1A. With this combination there are an equal number of homopolymer and heteropolymer melts in the fit, giving equal weight to each.

12 Each cap must be removed one at a time, and the length must be varied (see Figure 1A).

